# Spatial organization of the mouse chromosomes influences the landscape of intrachromosomal exchange aberration breakpoints

**DOI:** 10.1101/852277

**Authors:** Y.A. Eidelman, I.V. Salnikov, S.V. Slanina, S.G. Andreev

## Abstract

How 3D chromosome organization affects chromosomal aberrations is an important unresolved question in cell and radiation biology. In interphase the chromosomes form territories where chromatin folds into quite heterogeneous states. The mechanisms determining the spectrum of chromosome conformations remain poorly understood. We introduced the polymer model of mouse chromosome and generated the ensemble of 3D conformations. The chromosome model was validated against independent Hi-C (high-throughput chromosome conformation capture) data. The model described well Hi-C contact heatmap for chromosome 18 in pro-B mouse cells, both ATM-deficient and wild-type. We used the chromosome model to assess the role of chromosome structure in breakpoint distribution for intrachromosomal exchange aberrations. We investigated the effect of elevated frequency of breakpoints outside of the region of enzymatic breaksite. Chromosome aberration model explained breakpoint distribution under recurrent and ionizing radiation-induced DNA double-strand breaks in mouse chromosome 18 on the basis of contact-first mechanism. Overall, our results provide a framework for assessment of role of chromosome 3D organization on chromosome aberrations following DNA damage of different origin.

## Introduction

Exposure to ionizing radiation (IR) induces DNA double-strand breaks (DSBs) in the genome, and misrejoining of DSBs results in exchange-type chromosomal aberrations (CAs). Recurrent chromosomal aberrations are hallmarks of various cancers [1]. The probability of exchange events between damaged chromosomal sites or DSBs is likely to depend on the proximity of respective chromosomal loci. Hence the spatial organization of interphase chromosomes and the whole genome might be important prerequisite for radiation induced CAs and predispose some chromosomes to be targeted specifically resulting in cancer aberrations [2, 3]. However, the extent to which the spatial organization of chromosomes contributes to CAs is still unclear. To elucidate this connection, complex investigations are performed now both experimental and theoretical, including computational modeling, to study structure of interphase chromosomes and mechanisms of CA formation.

Reconstruction of spatial organization of interphase chromosomes is a complex problem of polymer physics [4] which is now becoming a hot topic of theoretical research in the field of genome biology [5–7]. There were several approaches to biophysical modeling of CA formation with chromosome structure taken into account [8–13].

They however met the serious problem, as to impossibility of well grounded 3D structure modeling due to lacking of comprehensive experimental data. The important role of chromosomal contacts in intra- and inter-chromosomal CA formation was pointed out [8]. Experimental assessment of contact frequencies between genomic loci became possible after development of Chromosome Conformation Capture technique and its whole-genome derivation Hi-C described in seminal papers [14, 15].

Hi-C study of human cells has revealed that human genome is folded according to a fractal globule model [15]. How the physical principles of chromosome organization are manifested in CA formation remains unclear. Investigation of chromosome folding is important for obtaining detailed information on chromatin/chromosomal proximity which impacts CA induction by IR and other factors.

The new possibilities to detect CAs with high precision arose when deep sequencing based technique HTGTS (High Throughput Genomic Translocation Sequencing) was developed [16]. This approach allows to determine genomic position of misrejoined loci or breakpoints along the chromosome with nucleotide-scale precision. In [17] complex influence of genome organization on translocation frequencies was assessed for G1-arrested mouse pro-B cell line by parallel characterization of 3D genome structure with Hi-C and breakpoint measurement with HTGTS. CAs were triggered by DNA DSBs of different origin, I-SceI endonuclease induced recurrent DSBs and IR (γ-rays) induced DSBs in the same cellular system.

To assess the role of chromosome structure in breakpoint distribution for intrachromosomal exchange aberrations we developed here the polymer models of mouse chromosome 18 in normal and ATM-deficient pro-B cells. The models well described the available mouse Hi-C data. On this basis we calculated frequencies of breakpoints from DSBs of different origin and their distribution along the chromosome. Our results provide new insights into complex interconnection between spatial and genomic separations between DSBs.

## Results

### Polymer model reveals spatial organization of interphase chromosome 18 in pro-B mouse cells

The polymer model of mouse chromosomes enabled us to investigate spatial organization of chromosome 18 in G1-arrested ATM-deficient (ATM^−/−^) and wild-type (WT) mouse pro-B cells. The model generated the statistical ensemble of chromosome conformations representing cell-by-cell variation of 3D structures. The model reproduced the whole chromosome Hi-C contact maps with high accuracy (Figure 1 a, b). The agreement between data and simulations is close on several Mbp scale as well (the example in the inset in Figure 1 a). Our method predicted the plaid pattern of contact maps for both cell lines without *a priori* information about compartments, unlike other approaches [4, 18]. Besides the whole map, the polymer model reproduced quantitatively pseudo-4C data, i.e. the segment of Hi-C map *P(i, j)* with fixed *i* (bin 70.6–70.7 Mbp), Figure 1 c, d. Comparison between two cell lines demonstrates that pseudo-4C data are visibly different, especially around the bait. The conformations of chromosome 18 in both cell lines are compact, heterogeneous globular structures. The examples of conformations, two from each ensemble, are shown in Figure 1 e, f.

**Figure 1.**
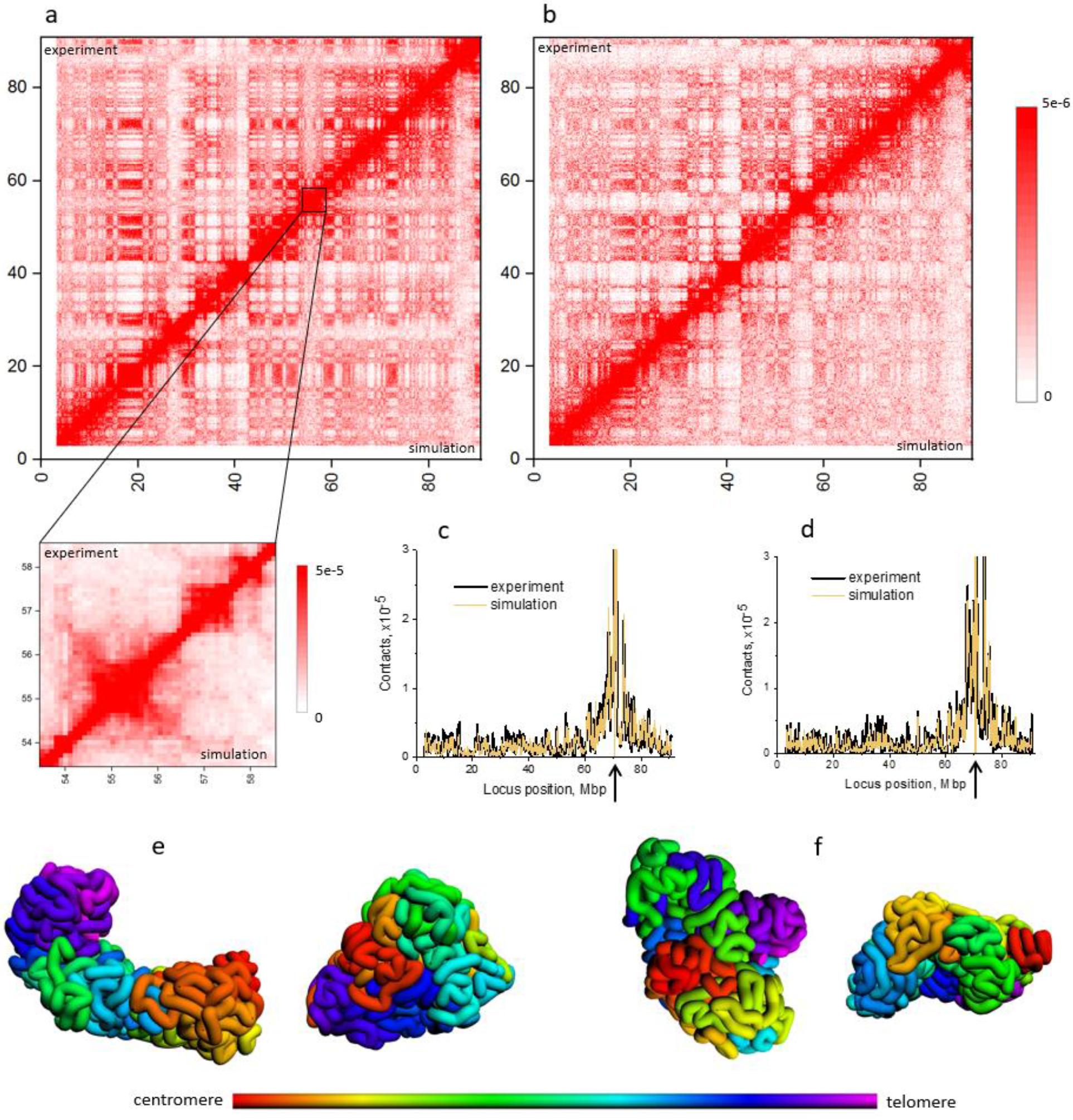
Globular organization of mouse chromosome 18. a, b: Hi-C maps of chromosome 18, ATM^−/−^ and WT pro-B cells, resolution 100 kbp. Upper triangle: experiment [17]; lower triangle: simulation. a: ATM^−/−^, Pearson correlation R=0.968. The inset shows quality of description on the lower scale, region 53.5-58.5 Mbp. R=0.970. b: WT, R=0.951. Between ATM^−/−^ and WT R=0.902. c, d: Pseudo-4C extracted from Hi-C data. Bait position: 70.6-70.7 Mbp (I-SceI locus), marked by the arrow. c: ATM^−/−^, Pearson correlation R=0.949; d: WT, R=0.937. Between ATM^−/−^ and WT R=0.916. e, f: Typical simulated conformations of mouse chromosome 18. e: ATM^−/−^, f: WT. Coloring indicates locus positions along the chromosome, see ideogram below.

In [15] the analysis of Hi-C data on human cells resulted in conclusion that chromosome structure in cell nucleus at the megabase scale is consistent with a fractal polymer globule model. For mouse cells the analysis of dependence between number of Hi-C contacts for pairs of loci and their genomic separation for the whole genome also showed that this dependence obeys the law ~*s*^−1.05^ for genomic separations 0.5-5 Mbp [17], that reveals the concept of genome fractality similar to human cells. However, the data averaged over the whole genome are not informative regarding individual chromosomes. To avoid this uncertainty we built several different functions, simulated as well as experimental, for chromosome 18 in ATM^−/−^ and WT cells. The first one is frequency of contacts at genomic separation *s* (*f*(*s*)), Figure 2 a, b. The sum of *f*(*s*) over all s is total frequency of contacts in the chromosome. Second, dependence between number of Hi-C contacts for pairs of loci and their genomic separation (Figure 2 c, d), the same as in [17] with the only difference that we built it for individual chromosome 18. Third, averaging of this dependence over all pairs of loci with given s, which is equivalent to function *f*(*s*)/(*N-s*) (Figure S1). All of these functions were analyzed for power law dependence.

**Figure 2.**
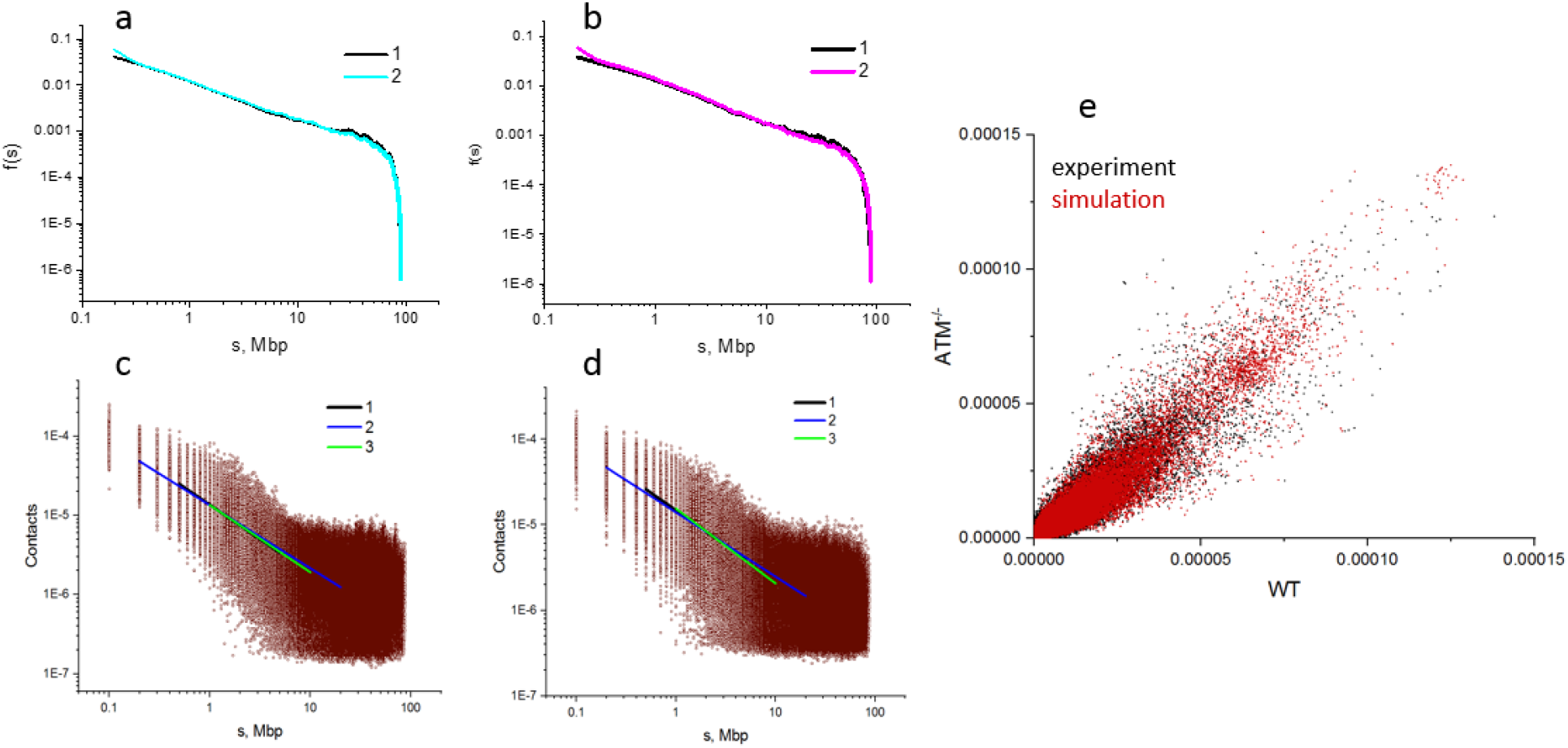
Structural features of mouse chromosome 18 in ATM^−/−^ and WT cells. a, b: contact frequency as a function of genomic distance. All experimental and simulated dependencies are described by the single function f(s)~s^k^ in wide range of s, 0.2 to 50 Mbp. a: ATM^−/−^. 1 – experiment [17], 2 – simulation. k=−0.89. b: WT. 1 – experiment [17], 2 – simulation. k=-0.88. c, d: Hi-C contacts vs genomic distance. Experiment [17], brown circles. The dependencies are fitted by function ~s^k^ with k varied for different ranges of s (colored lines). c: ATM^−/−^. 1 – range 0.5−5 Mbp, k= – 0.87. 2 – range 0.2−20 Mbp, k=−0.79. 3 – range 1−10 Mbp, k=−0.85. d: WT. 1 – range 0.5−5 Mbp, k=−0.82. 2 − – range 0.2−20 Mbp, k=−0.75. 3 – range 1−10 Mbp, k=−0.87. e: correlation between Hi-C contacts in ATM^−/−^ and WT cells. Black points – experiment, red – simulation.

Functions *f*(*s*) are described by the single power law in very wide range of *s*, 0.2 to ~50 Mbp. The slopes for two cell lines are close and equal ~-0.9 (Figure 2 a, b). The other functions, Figure 3c, d and Figure S1, are not described by the single power law in such a wide range, and in more narrow ranges their slopes vary from −0.75 to −0.87. Our analysis of all functions in various ranges of genomic separations built for both cell lines showed that slope close to −1 was never observed. The structure predicted by our polymer model and supported by mouse chromosome 18 Hi-C data is a nonfractal heterogeneous globule.

**Figure 3.**
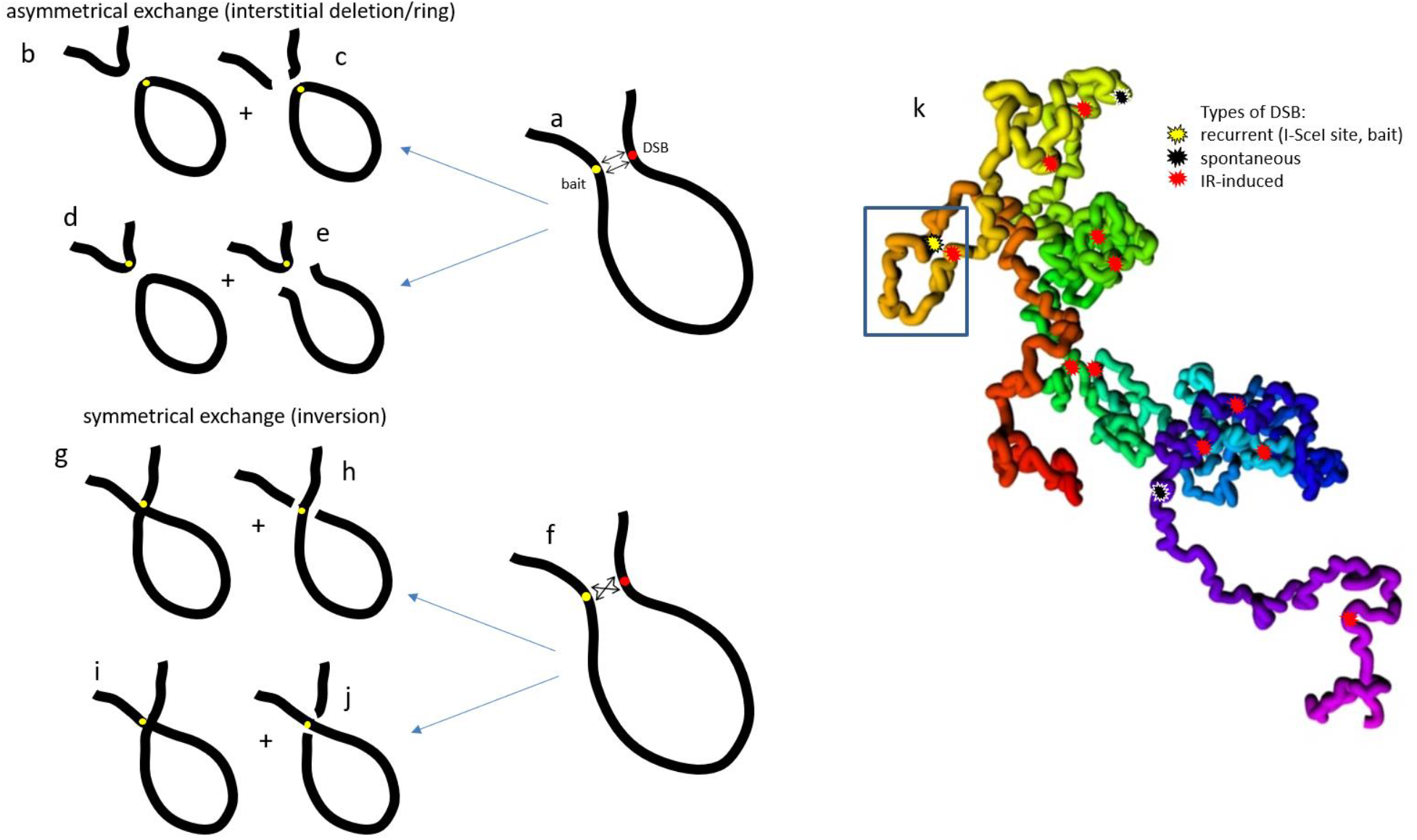
Intrachromosomal exchange aberration model. a-e: asymmetrical exchanges – interstitial deletions, rings. f-j: symmetrical exchanges – inversions. b, d, g, i: complete exchanges. c, e, h, j: incomplete exchanges. c: ring and broken chromosome, e: linear interstitial fragment, h, j: incomplete inversions. Different location of bait relative to rearranged segment: b, c, g, h – the bait inside rearranged segment; d, e, i, j – bait outside of rearranged segment. k: Intrachromosomal aberrations in simulated 3D structure of mouse chromosome. If the bait and another DSB are in contact (rectangle) they form an exchange with probability *P_c-e_*.

Comparison of chromosome 18 contact maps in two cell lines shows their high correlation (Pearson’s R=0.902, Figure 2 e), though some visual differences in contact patterns are seen (Figure 1 a, b). The origin of these minor differences could be due to differences in gene expression [17, 19]. Two functions of contact frequency, *f*(*s*) and *f*(*s*)/(*N-s*), have the same slopes for both cell lines, Figure 2 and Figure S1. Thus, our analysis agrees with the conclusion [17] that deficiency in ATM gene which is involved in many cellular functions including DNA repair is not reflected significantly on 3D organization, at least for chromosome 18.

### Model of intrachromosomal exchange aberrations

In [17] CAs were measured with HTGTS (High Throughput Genomic Translocation Sequencing) technique developed in [16]. In this technique the presence of an exchange aberrations involving the certain locus (bait) and location of the second locus is determined by sequencing chromosomal fragments containing the bait and their alignment to the reference genome.

To study the distribution of chromosomal rearrangements along the chromosome we used the following model and definition of rearrangements induced by DNA-damaging factors. The definition extends those used in [16]. Intrachromosomal aberrations in our model included following endpoints resulted from damage induction and misrejoining, Figure 3:

interstitial deletions in the form of rings and linear fragments, Figure 3 a-e;
inversions, Figure 3 f-j;

In our model CAs are rearrangements of mixed type. They are formed due to interaction of DSBs of different origin. I-SceI-induced recurrent DSB (bait) with IR-induced or spontaneous DSB (Figure 3) with different bait position, inside or outside of rearranged segment. On the chromosome itself the damaged loci rejoin and the points of junctions (breakpoints) are detected by HTGTS technique as exchange aberrations. The locations of all breakpoints are presented as distributions depending on breakpoint coordinate on the chain or on genomic separation from the site with I-SceI-induced DSB. Aberrations of bait-IR and bait-spontaneous origins were calculated separately. In the model we assume that DNA DSB does not result in the actual chromosomal break. Two DSB-bearing subunits enter the exchange process (misrejoining) as two entities rather than four free ends. In this regard our exchange model is consistent with Revell-type models [20]. This assumption is in concert with the observation of translocations *in vivo* [21]. In the literature there are alternative models of CA formation with incorporating movement of broken DNA free ends [10].

Experimental technique [16, 17] cannot distinguish between complete and incomplete exchanges. A complete exchange is an aberration where two participating DSBs misrejoin completely, i.e. form two junctions of DNA sequences located on both sides of both damaged loci, Figure 3 b,d,g,i. In an incomplete exchange aberration two DSBs rejoin partially, as a result only one junction is present, Figure 3 c,e,h,j. Experiment [17] keeps track only of junctions containing bait. In contrast, our model scores both complete and incomplete exchanges separately. Since we use only bait-containing junction for comparison with the experimental data, the information about presence or absence of the second junction is not used.

One can mention some limitations of both CA model and experimental data. Of all types of intrachromosomal joining events measured in [16, 17] we do not take into account only single-DSB events, resections, because they are on a very small scale, beyond our model’s resolution. Neither we nor [16, 17] score terminal deletions, since these are not DSB joining events.

In [16, 17] all aberrations measured are termed as “translocations”. This is not in line with the classical cytogenetical definition of translocations as symmetrical *inter*chromosomal exchange aberrations [22]. In order to avoid misinterpretation, hereafter we use term “intrachromosomal exchange aberrations” or “exchanges” rather than intrachromosomal “translocations” used in [16, 17], though the entities simulated are the same as measured in the experiments [17].

For the generated ensemble of chromosome conformations consistent with the experimental Hi-C data (Figure 1) we introduced the function of structural proximity Ψ_4c_(*s,x*), see also section “Chromosome aberrations landscape is shaped by damage proximity”. This function is the probability density of a chromosome subunit with genomic separation from the bait s being at spatial separation or distance *x* from bait. The bait was located in subunit 70.6-70.7 Mbp, in accordance to location of I-SceI site on chromosome 18 in [17].

To bridge two types of proximity information, on chromosomal structure and chromosomal damage, we introduced the new function of “damage proximity” φ_D,4c_(*s,x*), number of DSBs, or number of damaged, containing DSB, 100 kb chromosome subunits, with genomic separation s and spatial distance *x* from the bait. On the basis of this function for each subunit we calculated the frequency of contacts between DSBs in this subunit and DSB in the bait. Every DSB contacting with the bait forms the exchange, i.e. one or two junctions, with probability contact-exchange *P_c-e_*.

Since we deal with ATM-deficient cells, we do not take into account repair explicitly; the damaged subunits can only interact with each other, which is the meaning of *P_c-e_*. Possible movement of damaged subunits being not in contact initially is taken into account implicitly by introducing function *P_int_*(*x*), probability of interaction (misrejoining) of two damaged subunits separated by spatial distance *x* at the moment of irradiation.

Simulation of intrachromosomal aberrations was carried out by Monte Carlo technique, see Methods. The exchange was classified as symmetrical (inversion) or asymmetrical (interstitial deletion) with probabilities 0.5. The bait was placed in the rearranged (inverted/deleted) or in nonrearranged chromosome segment with probabilities 0.5. The exchange was incomplete (only one junction) with probability *P_inc_*. In most calculations we used *P_inc_*=0, i.e. no incomplete exchanges. For IR-induced aberrations fraction of incomplete exchanges is usually small, <5% [23]. Figure S2 shows the simulated dose-response curves for aberrations of various types and impact of non-zero incompleteness parameter *P_inc_*. Although aberrations are formed by two DSBs, or by two damaged subunits (at the doses considered damaged 100 kbp chromosome subunit may contain only one DSB, two and more are unlikely) the dose-dependences are linear. The reason is that one DSB (in the bait) is always present. The number of breakpoints as a function of locus position on chromosome was compared to the experimental data with resolutions 200 kbp and 2 Mbp.

### Relationship of intrachromosomal exchange aberrations with chromosome spatial organization

The model of chromosomal aberrations should score both spontaneous exchanges (i.e. with no radiation exposure) and those arising from interaction between IR-induced and I-SceI-induced DSBs. At the first stage we concentrated on simulation of spontaneous aberrations. For the ensemble of structures accurately describing the Hi-C data which was generated previously we found the distributions of spontaneous DSBs along the chromosome which gave rise to control breakpoint distributions fitting the experimental data.

At the next stage, we would compare our simulation results for IR-induced aberrations with the experimental data with resolution 200 kbp. Thus, control breakpoints needed also to be simulated with this resolution which is higher than 2 Mbp given for control presented in [17]. Besides, we took into consideration the uncertainties by reconstructing control with two scenarios (see Supplementary Methods). Both scenarios gave non-uniform distribution of DSBs (Figure S3 a, b) which resulted in quantitative description of the distributions of breakpoints in control with both resolutions, 200 kbp and 2 Mbp (Figure 4).

**Figure 4.**
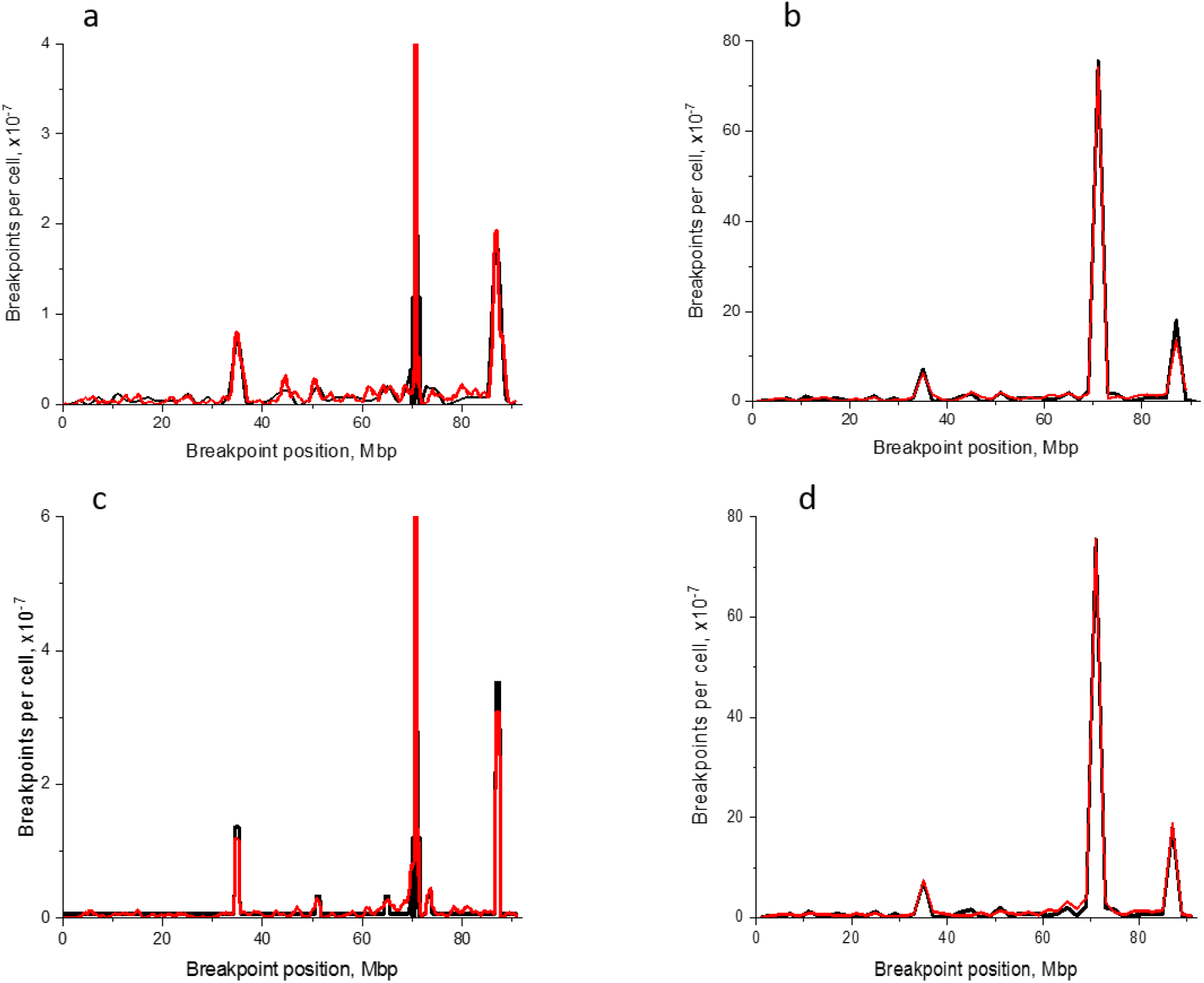
Reconstruction of background breakpoint distribution with higher resolution (200 kb). a, b: scenario 1. a: resolution 200 kbp. 1 – derived from [17], 2 – simulation. Pearson correlation R=0.898. Correlation was calculated for all points except the region within 1 Mbp from the bait. b: resolution 2 Mbp. 1 – [17], 2 – simulation. R=0.964. Correlation was calculated for all points except the bait-containing bin. c, d: scenario 2. a: resolution 200 kbp. 1 – derived from [17], 2 – simulation. Pearson correlation R=0.992. b resolution 2 Mbp. 1 – [17], 2 – simulation. R=0.979. Contact-exchange probability P=0.0029. Spontaneous DSB distributions – see Figure S3.

Fitting of the experimental distribution of breakpoints following 5 Gy *γ*-irradiation by our model for heterogeneous globular structure of chromosome 18 is shown with resolution 200 kbp and 2 Mbp in Figure 5 a, b. The control here was simulated by scenario 1. The similar result was obtained with control simulation by scenario 2, Figure S4 a, b. As the results for two scenarios of control simulation differ marginally with both resolutions, all further calculations were performed for scenario 1. Highly correlated description of the experimental breakpoint distribution (Figure 5, Figure S4) allowed us to determine contact-exchange probability *P_c-e_* which amounted to 0.0029.

**Figure 5.**
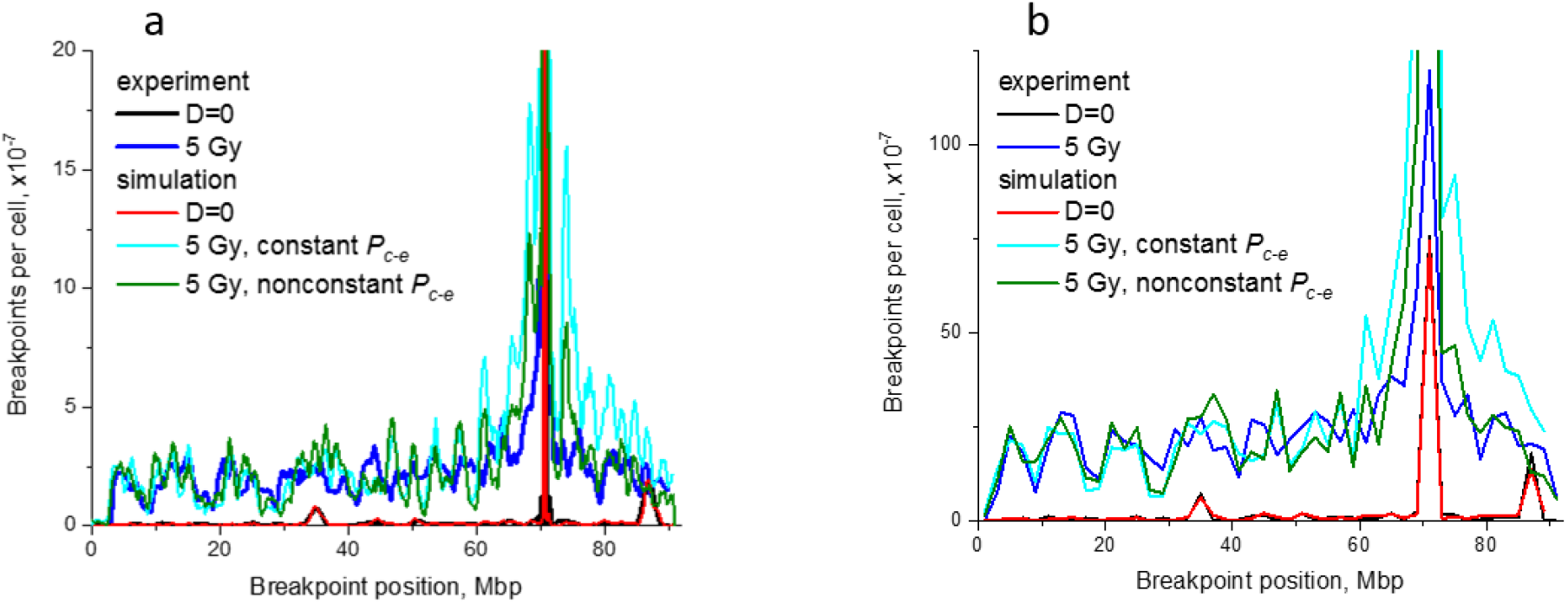
Heterogeneous globular model of chromosome 18 describes the experimental data on IR-induced breakpoint distribution. Control modelled by scenario 1 (Supplementary Methods). In both panels “experiment” refers to data [17]. Constant *P_c-e_*=0.0029. Nonconstant *P_c-e_*=0.0029 for j < 60 Mbp, 0.0017 for j≥60 Mbp. a: resolution 200 kbp. Pearson correlation between simulation and experiment for 5 Gy R=0.724 for constant *P_c-e_*, 0.732 for nonconstant. b: resolution 2 Mbp. Pearson correlation for 5 Gy R=0.781 and 0.789 respectively.

From Figure 5 and Figure S4 one can see that “distal” breakpoints (located before ~60 Mbp on chromosome 18) are described well with our CA model based on contact-first mechanism with the simplest assumption of *P_c-e_* being constant on condition that the chromosome is a heterogeneous globule. In the region close to the I-SceI site the model overestimates breakpoint frequency. This suggests that in the 10-20 Mbp vicinity of the I-SceI site exchange aberration formation is impeded compared to more distal regions. The possible explanation is lowered dynamics of damaged chromatin loci or DSBs. In [17] CAs were measured two days postirradiation, i.e. not only those formed immediately after exposure were scored. Our interpretation suggests that contact-first mechanism dominates at long genomic separations from the enzymatic DSB and role of DSB dynamics increases at relatively short separations. The alternative explanation is that contact-exchange probability is not constant, decreases closer to the I-SceI site. The simple model calculation was carried out where *P_c-e_* was 60% of the main value, i.e. 0.0017, for all subunits located after 60 Mbp, which improved the quality of fitting (Figure 5).

To clarify the role of structure, including compaction and structural heterogeneity, we simulated breakpoint distributions in control and after 5 Gy IR exposure for structures different from heterogeneous globule, homogeneous structures of chromosome 18: polymer coil, dense globule (with the same mean number of contacts as the heterogeneous globule) and more loose globule. The characteristics of the structures used are presented in Figure S5 e-g and in Table S1. The parameters of CA model for all structures are the same as for heterogeneous globule which agrees with Hi-C data. The resulting distributions with resolution 200 kbp are shown in Figure 6 a-d, with resolution 2 Mbp in Figure S5 a-d.

**Figure 6.**
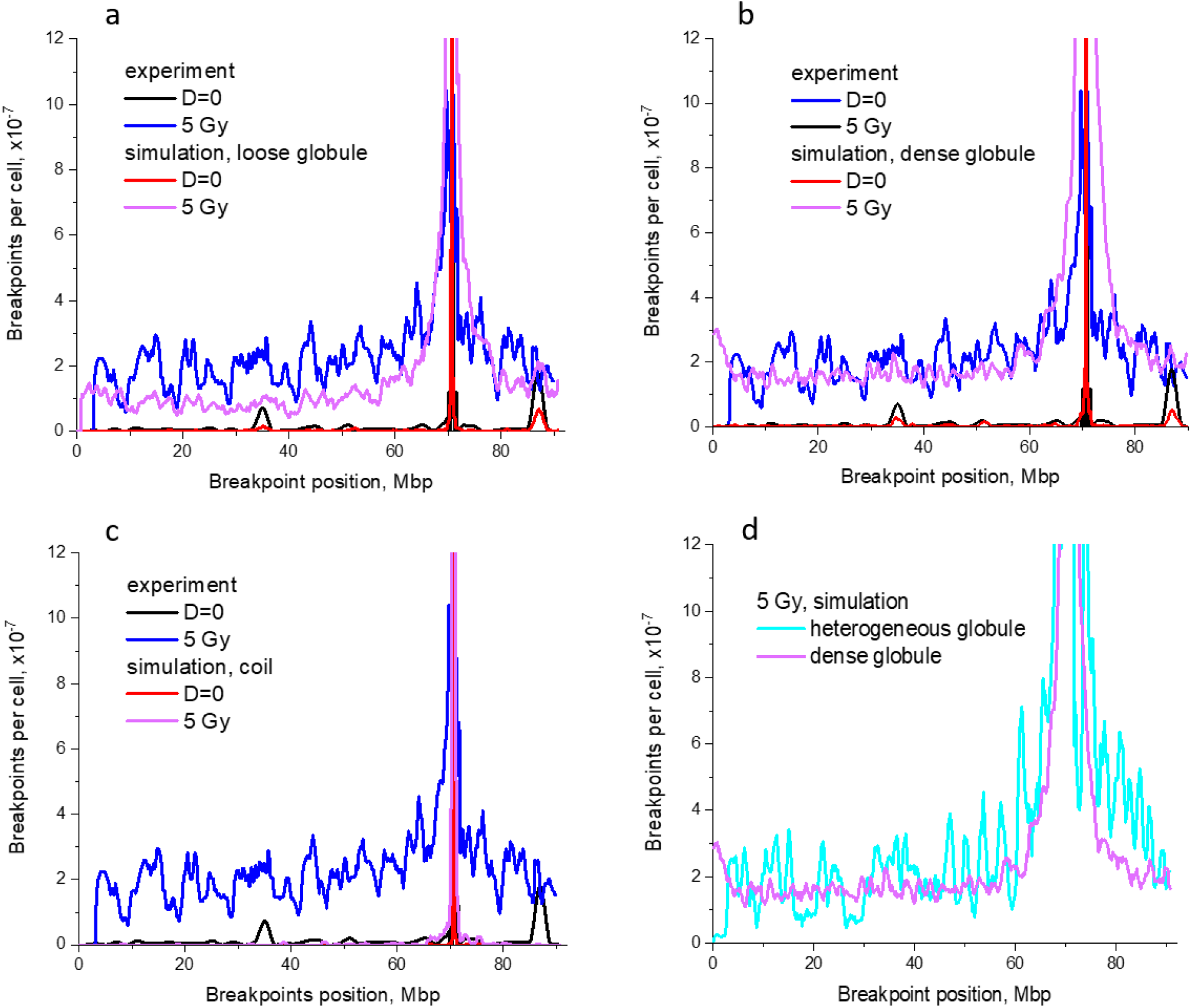
Role of large-scale chromosome structure in intrachromosomal breakpoint distribution of intrachromosomal exchange aberrations. a-c: different homogeneous structures of chromosome 18 vs experiment [17]. a: Pearson correlation between simulation and experiment for 5 Gy R=0.740. b: R=0.720. c: R=0.586. d: comparison between heterogeneous and dense homogeneous globules. In all panels resolution 200 kbp, contact-exchange probability *P_c-e_*=0.0029.

The globular structures describe the experimental data with similar quality, basing on Pearson’s correlation, for loose globule correlation is a little higher (0.74 vs 0.72 for dense globule and 0. 732 for heterogeneous globule). Homopolymer globule with the same degree of condensation as heterogeneous globule describes the *level* of breakpoints after 5 Gy irradiation fairly well but loses to heterogeneous globule in precision of peaks fitting. This suggests that the peaks on breakpoint distribution, apart from the main peak at bait, are related to heterogeneity of globular structure and contact map. In other words, the degree of globule heterogeneity determines fine behavior of breakpoint curve, its oscillations, especially in range 0-60 Mbp. On the other hand, the general structure of chromosome, its degree of compactization impacts dramatically the common shape of the breakpoint distribution: the peak at bait and relatively low-sloped decline in region up to ~10 Mbp from bait; the latter gives way to extremely steep decline if the chromosome is decondensed, i.e. after structural transition globule-coil (Figure 6 c). Presence of a very narrow (~1-2 Mbp) peak at bait for polymer coil is a result of increased frequency of local contacts even in a decondensed structure. The peak arises from local spontaneous interactions between subunits of polymer chain (on the conformation of coil one can see local clumps, Figure S5 f). In coil these interactions are much scarcer than in globule, hence the peak at bait is more narrow. The alternative explanation of peaks on breakpoint distribution is that contact-exchange probability (misrepair) is not constant, as in our simulation, and depends on genomic separation between enzymatic DSB site and IR-induced DSB. The other possibility may be that the repair rate depends on genomic separation [24].

### Breakpoint distribution following irradiation with different doses

After validation of the model by describing breakpoint data [17] for doses 0 and 5 Gy we calculated CA frequencies and breakpoint distributions in the wide range of doses, 0.1-15 Gy, to explore the impact of dose on the shape of breakpoint distributions. The distributions are presented in Figure 7 a, b with resolutions 200 kbp and 2 Mbp. The model predicts unequivocally elevation of breakpoint frequency with dose increase, > 5 Gy, in the wide range of genomic separations far from the bait. On the other hand, the same graph shoes that the model predicts decrease of breakpoints in the same range with dose decrease, < 5 Gy. This fact is explained in the model by the simple physical argument. Due to the condensed chromosome state any locus of chromosome contacts with the bait relatively frequently. On the other hand, due to uncorrelated DSB induction by γ-rays the probability of DSB induction in any subunit located far from the bait is high. Thus, the effect of significant increase of breakpoint frequency along the whole chromosome 18, including regions far from I-SceI site [17] is explained in the model by randomness of IR-induced DSB induction in the chromosome and its high degree of compactness due to folding in globule.

**Figure 7.**
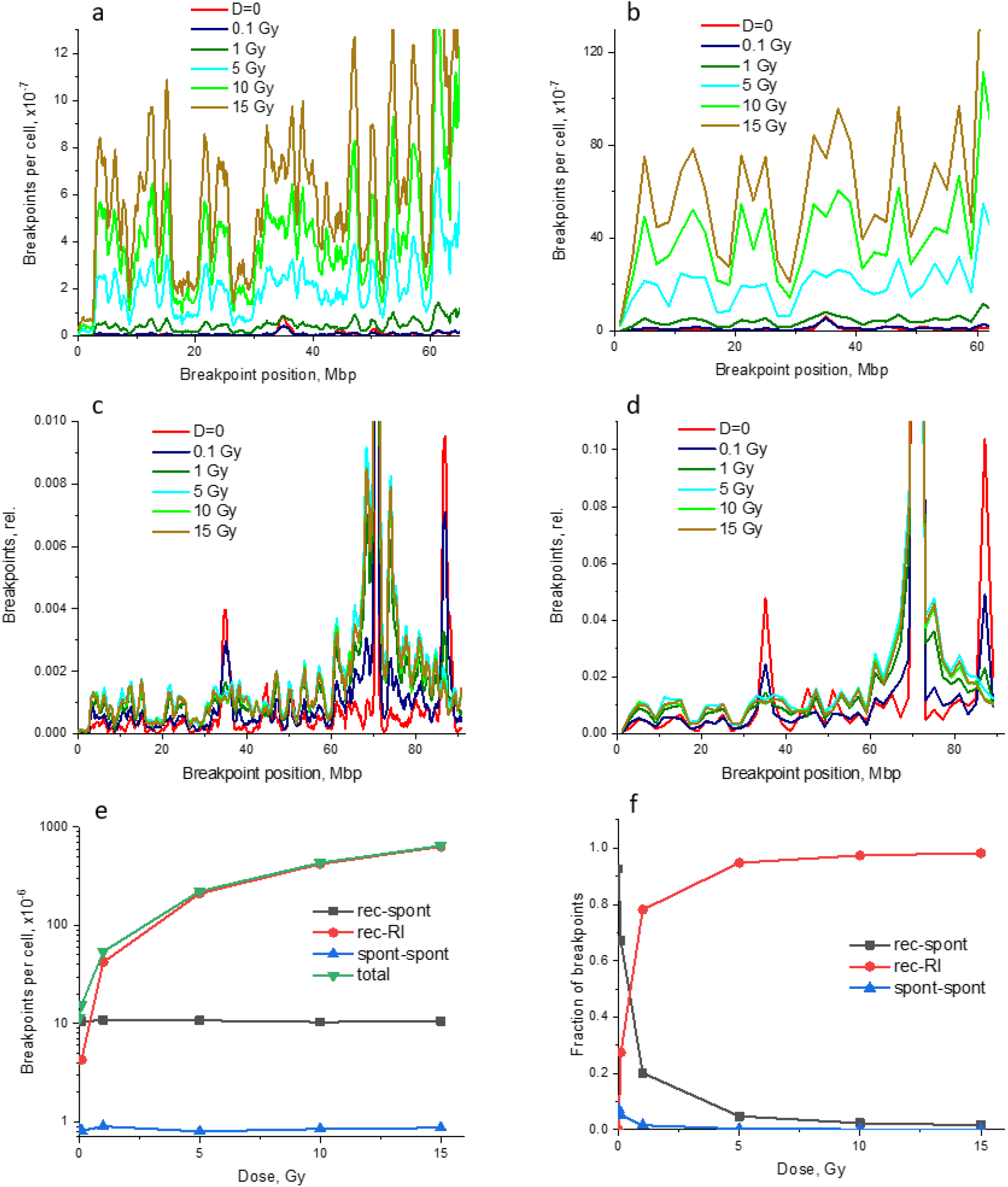
Breakpoints in the heterogeneous globular chromosome 18 at different doses. a, b: absolute frequencies of breakpoints in range 0-62 Mbp. a: resolution 200 kbp; b: resolution 2 Mbp. c, d: relative frequencies, normalized by the total number of breakpoints at given dose, in the whole chromosome. c: resolution 200 kbp; d: resolution 2 Mbp. Simulations for heterogeneous globule with 200 kbp resolution. e: dose dependency for absolute frequency of breakpoints, i.e. sum of breakpoint numbers per cell over all locus pairs, of different origin. Abbreviations: rec – recurrent DSBs; spont – spontaneous DSBs; RI – radiation induced DSBs. f: dose dependency for fraction of breakpoints of different origin. Abbreviations are the same as in panel e.

Since the induced DSBs are of different origin (recurrent DSB in the bait, spontaneous and IR-induced DSBs), the CAs formed have a mixed nature, not only recurrent (i.e. arisen from recurrent DSBs). To clarify which factors govern the shape of breakpoint distributions and to determine what is their contribution at different doses, we plotted the same dependencies normalized by unity (at each dose) with different resolutions, Figure 7 c,d. At low dose (0.1 Gy) the shape of breakpoint distribution is characterized by peaks at loci of high induction of spontaneous DSBs and, correspondingly, high spontaneous exchange aberration yields. Thus, at low doses spontaneous DSBs govern the shape of relative breakpoint curve.

The shape of breakpoint distribution after high doses of IR is dramatically different from distribution of enzymatic and spontaneous DSBs and their rearrangements. Misrejoining of IR-induced DSBs with I-SceI-induced DSB is the main factor determining breakpoint curve shape in ATM^−/−^ cells. This result (Figure 7 c,d) suggests (see also the previous section) that the mechanism of chromosomal rearrangements at pre-existing contacts of chromosomal loci (contact-first mechanism) correlates fairly well with yield of aberrations measured 2 days postirradiation. I.e. for chromosome 18 at least for long genomic separations from the bait contact-first mechanism of CA formation prevails, and subsequent dynamics of damaged chromosomal loci (in simple interpretation, DSB dynamics) towards each other, supposedly for repair, plays a minor role. Therefore, almost any 100 kbp chromosomal locus damaged by γ-rays can serve as a target for rearrangements with the enzymatic DSB in the bait if happens to be in contact with it (30 nm between subunit boundaries in these calculations) due to specifics of chromosomal conformations or fraction of conformations with given contact in the ensemble corresponding to Hi-C contact map. Thus the shape of intrachromosomal exchange distribution along the chromosome is governed by dose and distribution of contacting nondamaged and damaged subunits for given structural state, heterogeneous globule (Figure 1 e,f). See also the next section.

As to the contribution of different types of DSBs (enzymatic in the bait, spontaneous, IR-induced) to yield of exchange aberrations, the model predicts the following pattern, Figure 7 e,f. At the lowest dose considered, 0.1 Gy, most exchanges are formed due to interactions between bait and spontaneous DSBs. At dose as high as 1 Gy exchanges with IR-induced DSBs prevail, and at higher doses this component dominates.

### Chromosome aberrations landscape is shaped by damage proximity

Different kinds of proximity are distinguished in our model, proximity or structural contacts, at the moment of irradiation (contact-first mechanism) and proximity as a result of DSB movement toward each other after irradiation (breakage-first mechanism). In any case, predicting type and frequency of CAs and position of breakpoints requires the information on spatial distribution of lesions, i.e. distribution of spatial distances between them. But the only structural information available for all pairs of loci is Hi-C map. The proximity of damaged chromosomal subunits leading to CA formation does not come down to structural proximity in Hi-C sense. The distribution of spatial distances between lesions which were not yet eliminated by repair process is important for CA formation. The general problem is how to combine these two kinds of proximity.

To address this problem, we introduced two new functions which corresponded to structural and damage proximity, respectively. The first function, Ψ_4C_(*s,x*), is the joint probability density of a chromosome subunit having genomic separation *s* and spatial distance *x* from the bait. Function Ψ in general case can be defined for any pairs of subunits, but now we are interested only in the pairs where one subunit is the bait. This is designated by index “4C” in function’s name. If bait position on chromosome 18 is *i* and other locus position is *j*, than *i* is fixed (it is the subunit 70.6-70.7 Mbp) and genomic separation *s*=|*i-j*|. By introducing this function we move over to the space of variables including both 4C information and spatial distance information.

Figure 8 a shows the simulated structural proximity function Ψ_4C_(*s,x*) in 4C-X space for mouse chromosome 18 as heterogeneous globule. The integral of Ψ_4C_(*s,x*)d^3^*x* over Hi-C contact volume, i.e. over the distance from 0 to *R_cont_*, gives distribution Ψ_4C_(*s*) which is frequency of contacts between the bait and loci with genomic separation s from it. This distribution was already shown in Figure 1 c, d. The step beyond variable s into 4C-X space allows to obtain the information about structure (function Ψ_4C_(*s,x*)) as well as to move over to proximities for chromosomal lesions.

**Figure 8.**
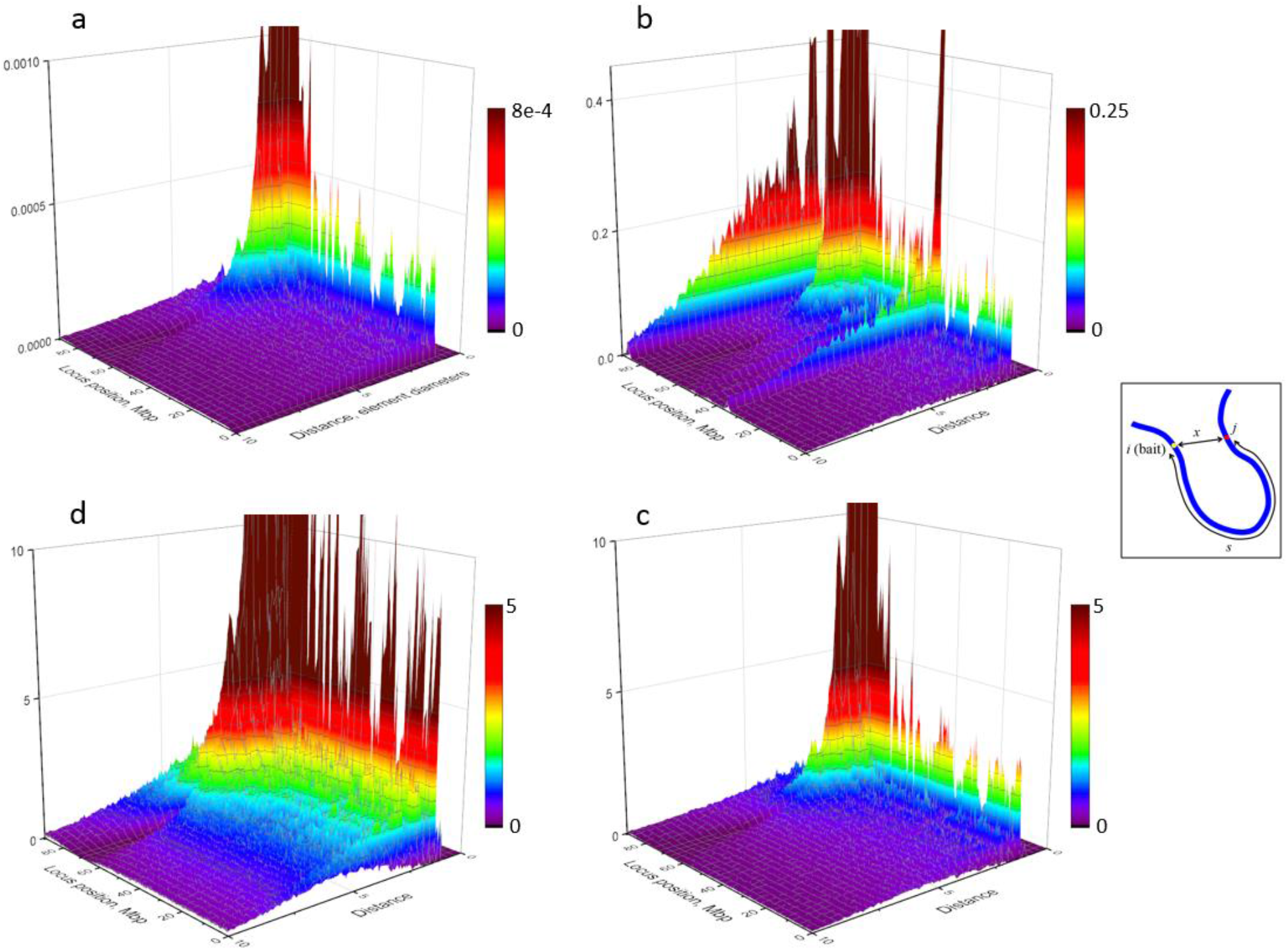
Structural and radiation damage proximity functions for mouse chromosome 18. a: structural proximity function Ψ_4C_(*s,x*) for bait position *i*=70.6-70.7 Mbp; *s*=|*i-j*|. Z scale is contact frequency. b-d: radiation damage proximity function φ_D,4C_(*s,x*) for different doses. b: D=0.1 Gy; c: D=5 Gy; d: D=15 Gy. Inset shows schematically genomic separation *s* and spatial distance *x* between the bait and locus *j*.

To go from purely structural proximity to proximity of chromosomal lesions we introduced function φ_D,4C_(*s,x*). It incorporates both structural and radiation dependencies because takes into account joint distribution of DSBs on chromosome over genomic separation s and spatial distance *x* from the bait, see Figure 8 b-d. The integral of φ_D,4C_(*s,x*)d^3^*x* over Hi-C contact volume gives distribution of DSBs with genomic separation s from the bait which are contacting with it (in Hi-C sense) and hence are prerequisite for CA formation by contact-first mechanism. This function provides the detailed information on contribution of lesions which shape breakpoint distribution, measured in experiment at given dose.

Figure 8 demonstrates that shape of functions Ψ_4C_(*s,x*) and φ_D,4C_(*s,x*) depends on dose and at low dose (0.1 Gy) they differ dramatically. Some degree of similarity of these functions at high doses is related to uniform distribution of IR-induced DSBs along the chromosome which is the case for sparsely ionizing radiation considered in the present work. In the case of exposure to densely ionizing radiation one can expect that structural proximity and lesion proximity would be different at doses higher than 0.1 Gy as well.

Function φ_D,4C_(*s,x*) (Figure S6 a) is the joint dependence of DSB frequency on two variables, genomic and spatial separations. From this function any partial dependence (on one variable with another fixed) can be extracted.

Figure S6 b,c shows examples of such distribution, slices of function φ_D,4C_(*s,x*). Depending on the place of the cut, these functions demonstrate different behavior which reveals complex relationships between along-the-chain proximity usually interesting to those who deals with Hi-C or 4C and spatial proximity reconstructed by our model.

All previous simulations were performed for contact-first mechanism of CA formation. This was expressed in maximal inter-DSB distance at the time of induction, at which they are able to interact, being the same as Hi-C contact distance. However, in the case of breakage-first mechanism this condition would not be imposed. To clarify how removal of restraint “lesion interaction distance = Hi-C contact distance” impacts intrachromosomal exchange aberrations, we considered more general case where damaged subunits not being in contact at the time of irradiation could interact too. To this end we replaced the contact-exchange probability with the function of interaction probability vs distance *x, P_int_*(*x*). In our model simulations, interaction probability is non-zero for distances between subunit edges up to 5x *R_cont_*. Besides, since particular shape of function *P_int_*(*x*) is unknown, we considered three significantly different shapes and determined its impact on breakpoint frequencies, Figure 9 a-f. CA formation from distances higher than Hi-C contact distance results in elevated breakpoint frequencies in all chromosomal loci compared to purely contact-first mechanism, Figure 9 d (on condition that function *P_int_*(*x*) is stepwise and the height of the step is the same as the value *P_c-e_* in previous simulations).

**Figure 9.**
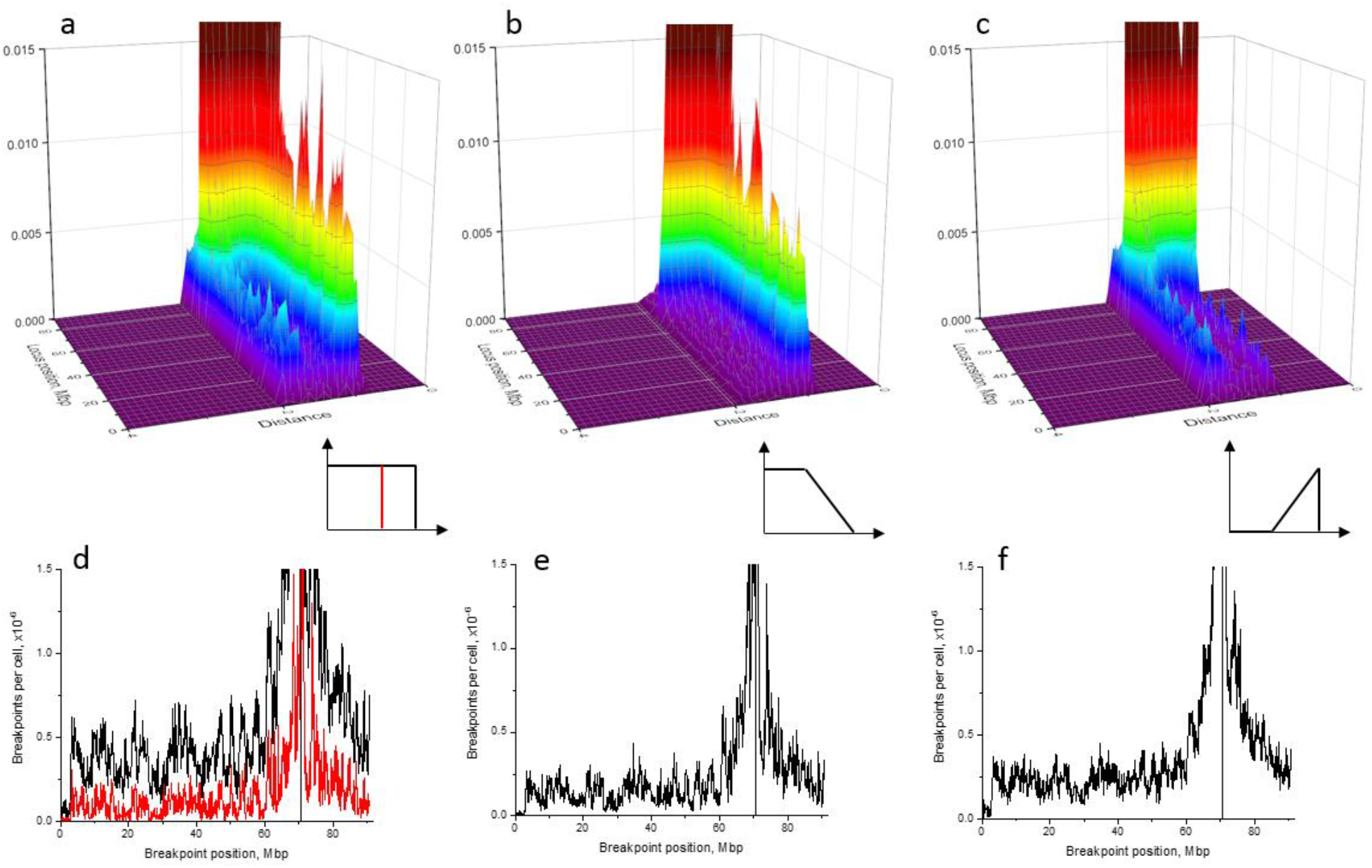
Interaction probability function impacts breakpoint distribution. a-c: product φ_D,4C_(*s,x*)·*P_int_*(*x*) for different *P_int_*(*x*). Corresponding *P_int_*(*x*) are shown in insets in lower right corners. In all cases dose=5 Gy, maximum interaction distance between subunit centers equals 2 diameters, maximum probability is 0.0029. Simulations for heterogeneous globular structure of chromosome 18. d-f: corresponding breakpoint distributions (resolution 100 kbp, no smoothing). Red line in panel d shows for comparison the distribution obtained for contact-first mechanism with the same interaction probability but maximal interaction distance equal to contact distance (red vertical line in the inset).

## Discussion

This work presents the first polymer model of the mouse chromosomes which generates cell-to-cell distribution of mouse chromosome 18 conformations and accurately reproduces the population Hi-C heatmap in both mutant (ATM deficient) and normal cells. The polymer model serves as the initial step for CA modeling on the basis of the “contact-first” mechanism, i.e. intrachromosomal exchange aberration formation at pre-existing contacts between subunits of chromosome.

The model predicts spatial organization of chromosome 18 in both cell lines as a heterogeneous globule with no sign of a fractal nonequilibrium state, which is associated with the inverse dependence of the contact frequency on genomic separation f(s)~s^−1^ at the megabase scale for human chromosomes [15]. Our conclusion differs from Hi-C analysis of mouse genome [17] revealing the fractal organization too. The difference in the principle of the chromosome organization may be explained by our use of a more accurate method for determining the slope of the frequency (or probability) of contact, not for the whole genome, i.e. with all chromosomes pooled, but for specific chromosome, and two different approaches to the calculation of the slope. The slope averaged over the genome used in [17] to characterize the type of organization of mouse chromosomes does not reflect the parameters of individual chromosomes. This question however requires further detailed consideration.

The potentials of volumetric interactions in polymer models of chromosome simulate attachment/detachment of various types of bridging molecules in polymer models of chromosomes [4, 5]. Our model describes Hi-C data not only at the whole chromosome level, but also at smaller scales, down to several TADs. At the same time, we do not take into account possible involvement of cohesin and extrusion mechanism in the TAD formation. Thus, at 100 kbp Hi-C resolution a role of cohesin and the extrusion in TAD modeling and heatmap reproducing is dispensable, unless we consider experiments with cohesin removal. Another important point is whether TAD structure contributes to the chromosome translocation formation and to what extent. There are no data in this regard and this question is subject for future consideration. Recent observations [25] shed new light on the mechanisms of DSB and γH2AX foci induction and repair within TADs. In the present model TADs participate in exchange aberrations as structural blocks within chromosome of increased frequency of contacts of 100 kbp bins or chromosomal elements. And this simple assumption is sufficient to describe CA data.

The CA model does not take into account the dynamics of DSBs or damaged chromosome subunits as only contact-first type mechanism is considered. Such dynamics can contribute to translocation yield [26] at different times after irradiation and may change the form of dose-response curves. At genomic distances about 10 Mbp from I-SceI site (bait) CA frequencies deviate from those predicted from chromosomal contacts. This may be an indication of diffusion of damaged chromosomal subunits contributing to the search for a partner for intrachromosomal interactions and misrejoining. The investigation of this issue will require further modification of our model to explicitly take into account the dynamics of loci, repair, *etc*, as well as experimental HTGTS data for normal cells. In addition to the hypothesis of the formation of CAs by pre-existing contacts, we assessed the case where CAs are formed from DSBs induced at distances greater than the HI-C contact size. This is still a rough assessment of the possible contribution of damage dynamics.

We show that heterogeneity of structure *via* the pattern of contacts, as well as globular state of chromosome are important contributors to the shape of CA breakpoints curve. The influence of heterogeneity is stronger at large genomic separations from the bait position (> ~10 Mpp apart), and weaker in closer regions (< ~10 Mpp). A decondensed chromosome state in the form of a polymer coil does not explain data on breakpoints after 5 Gy irradiation in the whole range of genomic separations. Thus, the globular state of the chromosome is necessary for the observed dramatic excess of breakpoints over the control level at large genomic separations from the sites of the recurrence of DSB, on the level of entire chromosome length. Unexpectedly, some aspects of the shape of breakpoint distribution are also consistent with a simple homogeneous globular chromosome model without heterogeneity of its internal contacts.

Hi-C does not measure 3D coordinates, only position of contacting subunits on a chromosome. For CA prediction, the distributions of chromosome subunits’ coordinates are important. Our model introduces joint genomic and spatial distance distribution for all pairs of DSBs or chromosome damaged sites. In this sense, we deal with different kind of proximity, structural and damage proximity, necessary for the predictions of the CAs. We show its complex shape in *s-x* space of variables which is dose dependent. Joint proximity distributions will be even more complex if high LET radiation (ions) is used as a source of DSBs in cellular DNA together with recurrent DSB or alone. This is due to the fact that for densely ionizing radiation the energy distributions in the particle tracks are correlated and the distribution of DSBs along the chromosome is quite different from uniform as for γ-rays.

An additional note about the difference between structural and damage proximity related to aberrations. Hi-C, like all other Chromosome Conformation Capture methods, measures spatial proximity in the sense of a contact being a formaldehyde crosslink between chromatin sites.

We estimate this Hi-C contact as a spatial distance of about 30 nm or less between the edges of the subunits, other estimations [27] are about 48 nm. At 60-90 nm separation, formaldehyde crosslinking of two chromatin loci is no longer possible even for large fixation time [28]. In this sense, the contact is short-range or “local” and differs from the range of the lesion interaction leading to CA, which is usually estimated as about 0.5 μm [9].

In conclusion, we developed the new methodology to predict CAs under exposure to physiological and environmental factors with accurate taking Hi-C structural information into account. We presented the first polymer model of spatial organization of mouse chromosome in ATM-deficient and WT pro-B cells. It generates the population of 3D structures which accurately captures pattern of chromosome Hi-C contacts with 100 kb resolution, available in [17]. The polymer model reveals that mouse chromosome 18 is organized in interphase as the heterogeneous polymer globule rather than the fractal nonequilibrium structure. We demonstrate the difference between structural proximity, measured by Hi-C technique, and chromosome lesion proximity, necessary for CA prediction. The interconnection between these two kinds of proximity is not measured by current experimental techniques. We reconstruct the hidden information by introducing the new probability function incorporating both structural and damage proximity and demonstrate its importance for analysis of intrachromosomal aberration breakpoints. Our model identifies the main factors shaping the CA and breakpoint landscape at different doses of ionizing radiation exposure. These findings create a novel framework to elucidate the mechanisms of CA formation with integration of 3D genomics and radiation biophysics information. This approach may have an important applications in predicting the therapeutic effects of radiation and chemotherapy on the chromosomes of cancer cells with the formation of various types of CAs, both stable and unstable.

## Methods

The model consists of two main sections: generating the ensemble of chromosome conformations with requirement of reproducing Hi-C data and simulation of intrachromosomal exchanges involving I-SceI site in control and following γ-irradiation.

The interphase structure of mouse chromosome 18 was modeled with the inhouse software using molecular dynamics package OpenMM [29, 30]. A chromosome was considered as a polymer chain of 908 subunits with 100 kbp DNA content. The interaction potential between any pair of non-neighbor subunits (*i, j*) consists of two components, the excluded volume potential and the attraction potential. The former is the same for all pairs of subunits, while the latter depends on the position of the subunits along the chain. This is ideologically similar to the method developed in [26] but applied to whole chromosome simulation. The coefficients for the attraction potential should be determined so that the ensemble of conformations generated by the model fits the experimental Hi-C map. To this end we developed the iterative algorithm described in Supplementary methods.

Next, the generated conformation ensemble was used for CA simulation by Monte Carlo technique. DSBs of different origin (I-SceI site, spontaneous, IR-induced) were distributed along the chromosome. IR-induced DSBs were distributed uniformly, spontaneous non-uniformly (for details of spontaneous DSB generation see Supplementary Methods), the bait always had 1 DSB (Figure 3 k). The chain subunit having a DSB is termed as “damaged subunit” and the further calculations were carried out in these terms. We neglect the probability of more than one DSB induced in a 100 kbp subunit which is reasonable at the doses of interest. If two damaged subunits were found to be in contact in the given conformation, they interacted, resulting in CA formation, with probability *P_c-e_*. The detailed description of the modelling procedure is provided in Supplementary Methods.

## Acknowledgements

The study was supported by MEPhI Academic Excellence Project (Contract No. 02.a03.21.0005); Russian Foundation for Basic Research grant 14-01-00825; The research is carried out using the equipment of the shared research facilities of HPC computing resources at Lomonosov Moscow State University and Joint Supercomputer Center of the Russian Academy of Sciences.

## Author Contributions

Y.E.: software, scientific analysis, work on manuscript. I.S.: software. S.S.: data analysis. S.A.: conceptualization, scientific analysis, work on manuscript.

## Conflict of Interests

The authors declare they have no conflicts of interest.

