## Supplementary information for "Spatial organization of the mouse chromosomes influences the landscape of intrachromosomal exchange aberration breakpoints"

### Contents

Supplementary Methods

Structure

Spontaneous aberrations

Radiation induced aberrations

Supplementary References

Supplementary Figures

Supplementary Tables

### Supplementary Methods

#### *Structure*

Mouse chromosome 18 was considered as a self-avoiding polymer chain of 908 subunits (monomers) with 100 kbp DNA content. This choice was due to the best available resolution of Hi-C maps available [1]. First 30 subunits (3 Mbp) correspond to the centromeric region. Subunit diameter was taken as 150 nm, but in most calculations spatial parameters are in units of monomer diameter, unless otherwise stated. The interaction potential between any pair of non-neighbor elements (*i, j*) separated in space by distance *r* consists of two components, $U_{ij}(r) = U_{ev}(r) + U_{attr}(r, i, j)$, where $U_{ev}(r)$ is an excluded volume potential, $U_{attr}(r, i, j)$ is an attraction potential depending on the position of the elements along the chain. The excluded volume between any two elements was modeled by the shifted Lennard-Jones potential with coefficient $U_{0ev}$. The attraction potential was modeled by the attracting component of the Lennard-Jones potential with a coefficient depending on the position of the elements along the chain.

To determine these coefficients, we developed an iterative algorithm consisting of the following steps:

1. Initial coefficients are equal for all element pairs, $U_0$.

2. The ensemble of up to 4000 conformations is simulated. The initial state of the chain is random walk with excluded volume, the system is equilibrated until gyration radius ceases to change.

3. The contact map with 100 kbp resolution is calculated by counting contacts between all pairs of elements (*i, j*) with genomic separation $s=|i-j|>1$ in all conformations in the ensemble generated. Two elements are deemed being in contact if distance between their centers is less than $R_{cont}$. Pearson's correlation between the simulated map and the experimental one is calculated. The experimental maps [1] were corrected for biases according to the coverage of each bin in the experiment [2]. For comparison both contact maps are normalized by total number of contacts. Again, only genomic separations $s>1$ are taken into account.

4. If Pearson's correlation between the contact maps ceases to improve, the simulation is stopped. Otherwise, for each element pair (*i, j*) the coefficient in the attraction potential increases if the simulated contact frequency is lower than the experimental one and decreases if it is higher.

5. Return to step 2.

Contacts with centromeric monomers are removed from consideration in steps 3 (contact map calculation and comparison) and 4 (modification of potentials), as the centromeric region is not assayed in Hi-C.

The algorithm described above was applied to WT and $ATM^{-/-}$ cells. The sufficient number of iterations was 15 and 13, respectively. For subsequent comparison with data on breakpoints involving I-SceI induced DSB, from the whole Hi-C maps only contacts with subunit (70.6-70.7 Mbp) were used, as I-SceI site on chromosome 18 in [1] is in locus 70.689 Mbp. This subset of Hi-C data is equivalent to 4C experiment with bait in this locus. The term "pseudo-4C" will be used hereafter.

***Spontaneous aberrations***

The distribution of spontaneous, i.e. unrelated to IR exposure, DSBs along the chromosome is unknown *a priori*. It should be selected so that the simulated distribution of breakpoints in the absence of IR agrees with the experimental one [1]. The problem is that the model's resolution is 100 kbp and the distribution in [1] is provided at 2 Mbp resolution except the small 2.5 Mbp region around the bait. We cannot know what is the share of each of 20 subunits comprising the given 2 Mbp bin.

Scenario 1. The experimental data with 2 Mbp bining [1] are interpolated by Akima spline with 100 kbp bin. The interpolated function is normalized so that total number of breakpoints equals initial one. Thus, in case of increased breakpoint frequency in given 2 Mbp bin this increase is "spread" over subunits comprising this bin. Next, in range 69.6-72.0 Mbp this distribution is replaced with the exact one, provided in [1] with 25 kbp resolution and grouped into 100 kbp bins. After that, we found the distribution of spontaneous DSBs (Figure S3 a) giving rise to breakpoint distribution which agree with that derived from the experiment (Figure 4 a).

Scenario 2. First, we determined average breakpoint frequency over all 2 Mbp bins except the peaks 34-36 Mbp and 86-88 Mbp and the region 68-72 Mbp. It amounts to 1.72.

Under hypothesis of uniform breakpoint distribution along the chromosome, their number in each bin should be distributed according to Poisson statistics with the mean 1.72. In this case probability of more than 3 breakpoints in a bin is 0.096, more than 4 breakpoints – 0.031. Therefore, all bins having more than 4 breakpoints are considered nonrandom with significance level 0.031, asterisks in Figure S3 c). In all other bins breakpoints are considered random with significance level 0.096 or higher.

We suppose that in each 100 kbp bin within a "random" 2 Mbp bin the mean number of BPs is 0.086 (then the mean number of breakpoints in 2 Mbp bin is 0.086x20=1.72). In "nonrandom" 2 Mbp bin with *n* breakpoints: all but one 100 kbp bins have 0.086 breakpoints, one bin has *n*-0.086x19 breakpoints. The choice of the 100 kbp bin with elevated number of breakpoints is arbitrary, we choose the bin in the middle of 2 Mbp bin. In region 69.6-72.0 Mbp, as in scenario 1, we use the exact experimental numbers of breakpoints.

Next, as in scenario 1, we found the distribution of spontaneous DSBs (Figure S3 b) giving rise to breakpoint distribution which agree with that derived from the experiment (Figure 4 c).

For both scenarios: in further simulations (of IR-induced breakpoints) theoretical distributions will be compared with the experimental one obtained with resolution 200 kbp and smoothed with sliding window

1 Mbp [1], thus we did the same procedure for control too. In region 69.6-72.0 Mbp the distributions were not smoothed.

To validate our scenarios, we compared the simulated control with the experimental data at 2 Mbp resolution (Figure 2 b,d). Both scenarios provide high level of agreement with data [1].

***Radiation induced aberrations***

In the case of irradiation with dose D we distribute both spontaneous and IR-induced DSBs along the chromosome. The distribution of IR-induced DSBs is uniform, the mean $8.2 \times 10^{-9}$ $Gy^{-1}$ $bp^{-1}$ [3]. Probability of induction of more than 1 DSB of any origin is neglected.

After determining number and positions of all DSBs the radiation proximity function $\varphi_{D,4C}(s,x)$ is calculated, number of DSBs with genomic separation *s* and spatial distance *x* from the bait. From this function the frequency of contacts between DSB in given subunit and DSB in the bait is determined as follows:

$$f_{D\ cont}(s) = \int_0^{R_{cont}} \varphi_{D\ 4C}(s,x) 4\pi x^2 dx \qquad (1)$$

Since every contact gives rise to exchange with probability $P_{c\text{-}e}$, the frequency of breakpoints in the subunit with genomic separation *s* from the bait is

$$f_{D\ BP}(s) = f_{D\ cont}(s) P_{c-e} \qquad (2)$$

This relationship was used for assessment of $P_{c\text{-}e}$. This value was chosen to fit qualitatively breakpoint level on the majority of chromosome 18, from 3 to ~60 Mbp (Fig.3). The resulting value is $P_{c\text{-}e}$=0.0029. In general case it can depend on locus position on the chromosome.

Breakpoint distributions are simulated with resolution 100 kbp (DNA content of a chromosomal subunit in the model). For comparison with the experimental distribution built with resolution 200 kbp and smoothed with 1 Mbp sliding window [1] we did the same procedure. As in control, the region 69.6-72.0 Mbp was not smoothed.

To take into account the possibility of interaction between DSBs being not in contact at the time of irradiation, we introduce function $P_{int}(x)$, probability of interaction between two damaged subunits vs their spatial distance *x*. For breakpoint frequency determination in this case equations (1) and (2) are replaced with

$$f_{D\ cont}(s) = \int_0^{R_{int}} \varphi_{D\ 4C}(s,x) P_{int}(x) 4\pi x^2 dx \qquad (3)$$

where $R_{int}$ is the maximal distance of interaction. The particular shape of function $P_{int}(x)$ is unknown. We considered three different model dependencies to clarify to what extent they impact breakpoint distribution for the same structure of the chromosome.

### Supplementary Figures

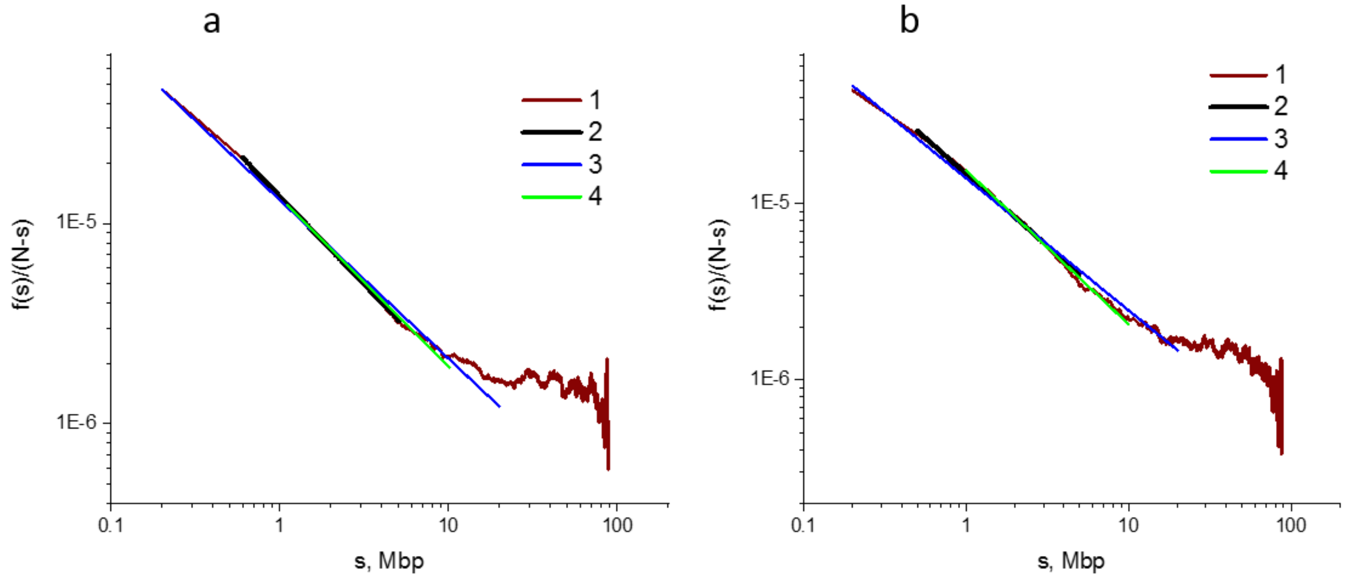


Figure S1. Renormalized contact frequency dependence on genomic separation. a: ATM$^{-/-}$, b: WT. 1 – function $f(s)/(N\text{-}s)$, equivalent to averaging contact frequencies for each $s$ separately. 2-4 – fitting of the data (curve 1) by function ~$s^k$. The ranges of $s$ and $k$ are the same as in Figure 3 c,d.

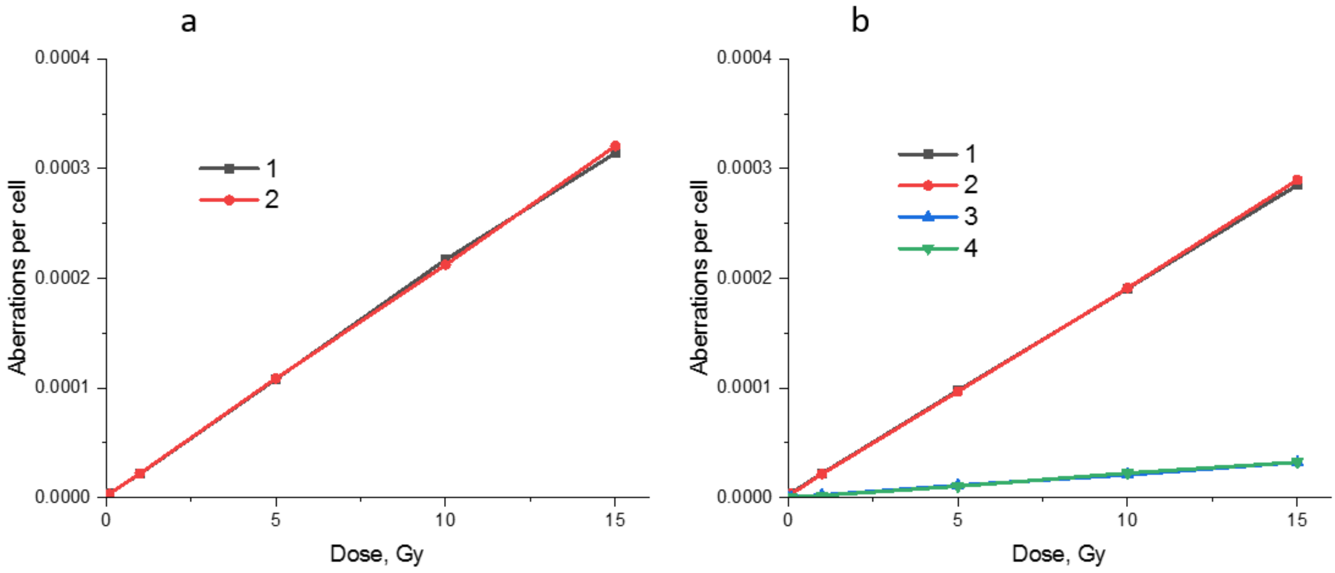


Figure S2. Dose-response curves for different types of intrachromosomal exchanges scored by the model. Simulation for chromosome 18 as heterogeneous globule, contact-first mechanism, $P_{c\text{-}e}$=0.0029. a: no incompleteness. 1 – rings, 2 – inversions. b: incomplete exchange probability $P_{inc}$=0.1. 1 – complete rings, 2 – complete inversions, 3 – incomplete rings and interstitial deletions, 4 – incomplete inversions.

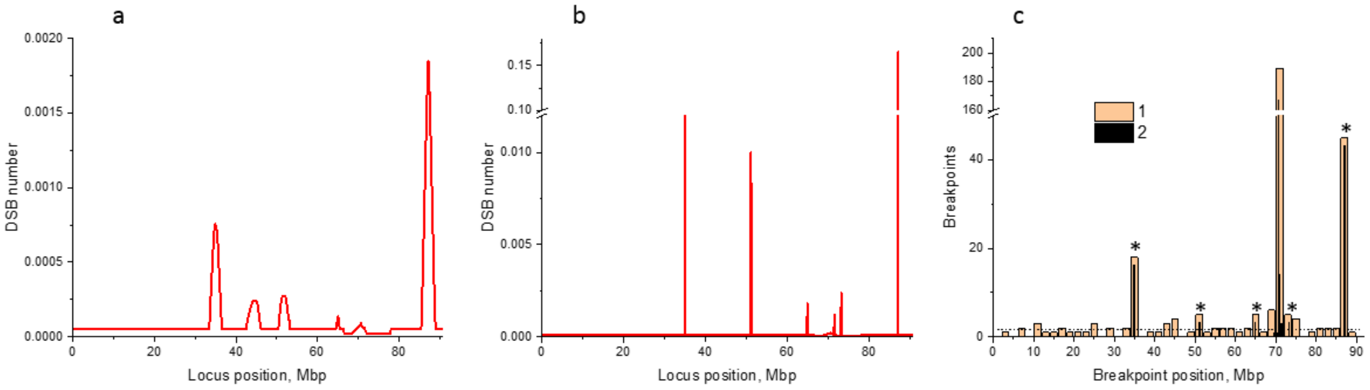


Figure S3. Reconstructed spontaneous DSB distribution along chromosome 18. The mean number of spontaneous DSBs in the 100kbp element reconstructed according to scenario 1 (a), according to scenario 2 (b). c: determining nonrandom breakpoint peaks in scenario 2. 1 – breakpoint distribution [17], resolution 2 Mbp. 2 – derived breakpoint distribution with resolution 100 kbp. Dashed line indicates the estimated level of random breakpoint distribution. Asterisks: loci with nonrandom breakpoint distribution, see text for details.

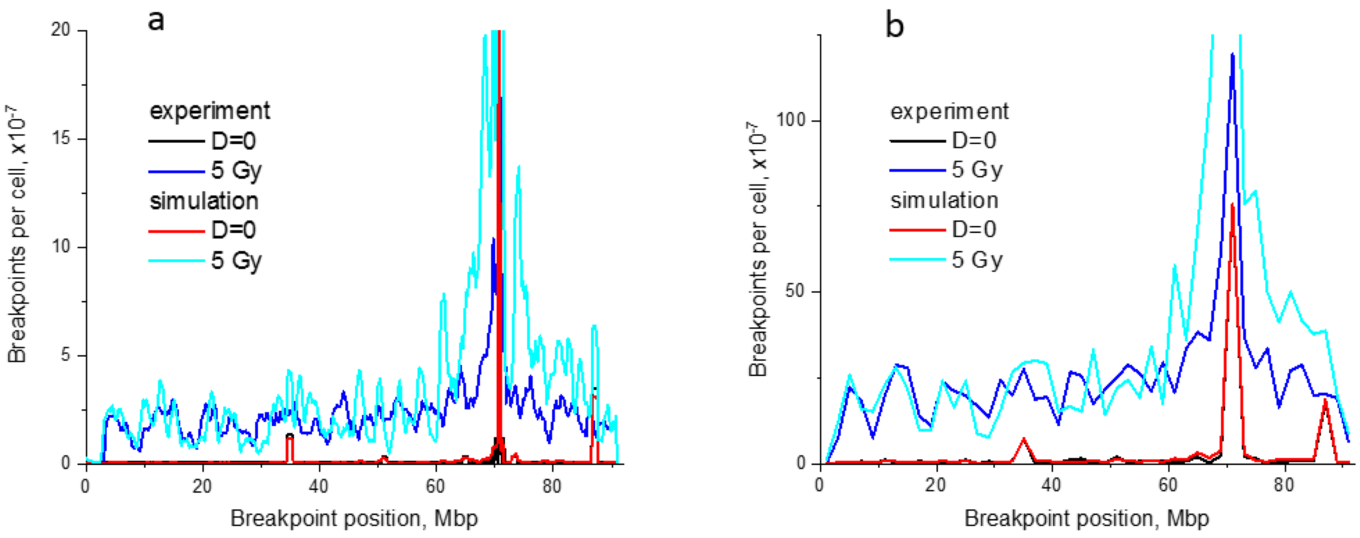


Figure S4. Heterogeneous globular model of chromosome 18 describes the experimental data on IR-induced breakpoint distribution. Control modelled by scenario 2. In both panels “experiment” refers to data [17]. a: resolution 200 kbp. Pearson correlation between simulation and experiment for 5 Gy R=0.722. b: resolution 2 Mbp. For 5 Gy R=0.786.

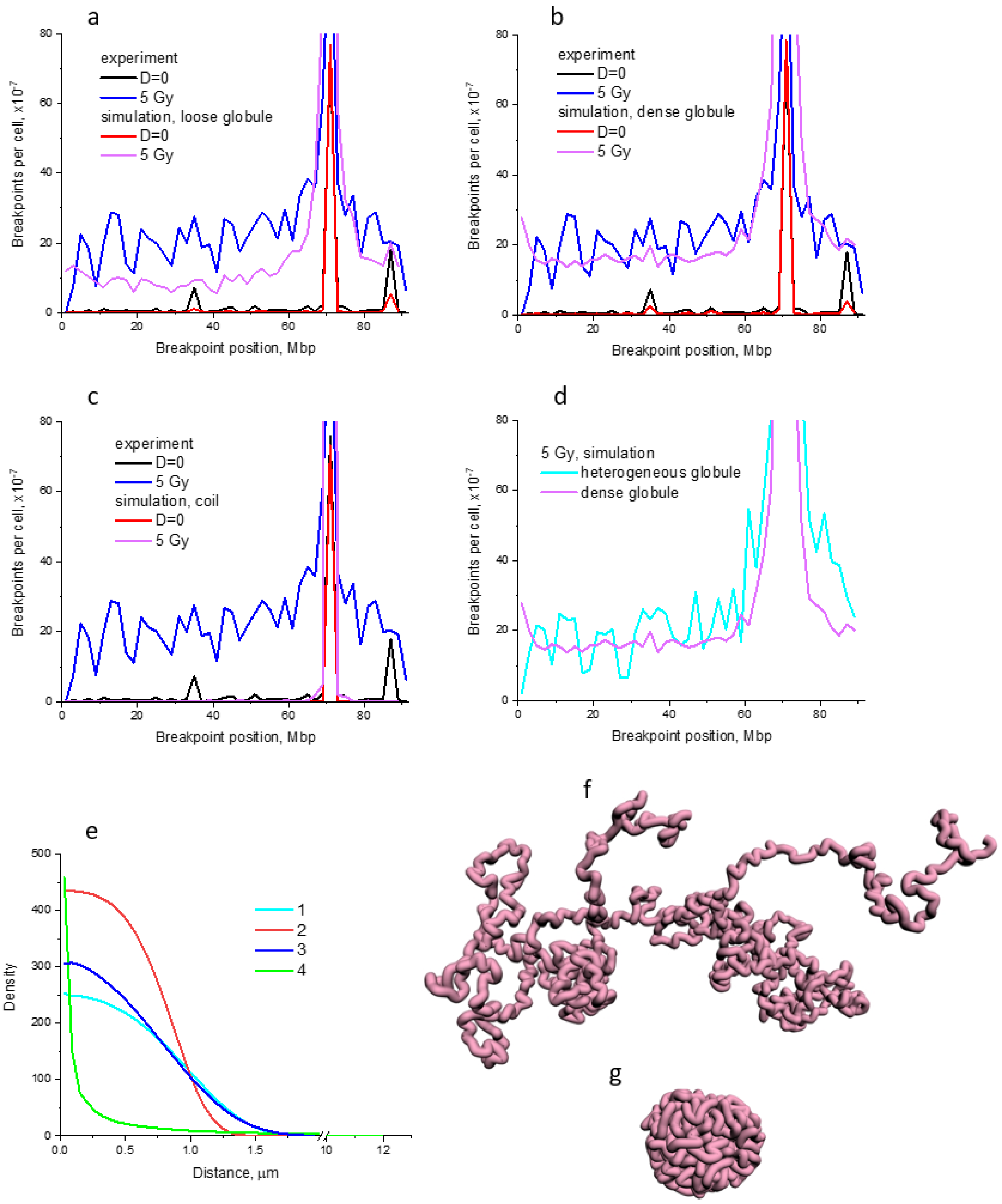


Figure S5. Role of large-scale chromosome structure in breakpoint distribution of intrachromosomal exchange aberrations. a-c: different structures of chromosome 18 vs experiment [17]. a: loose homogeneous globule. Pearson correlation between simulation and experiment for 5 Gy: R=0.742. b: dense homogeneous globule. R=0.748. c: coil. R=0.701. Resolution 2 Mbp, contact-exchange probability $P_{c\text{-}e}$ =0.0029. d: role of heterogeneity: comparison between heterogeneous and dense homogeneous globules. e-g: characteristics of structures used. e: radial density. 1 – heterogeneous globule. 2 – dense homogeneous globule. 3 – loose homogeneous globule. 4 – coil. f: the typical conformation of the polymer coil. g: the typical conformation of the dense homogeneous globule.

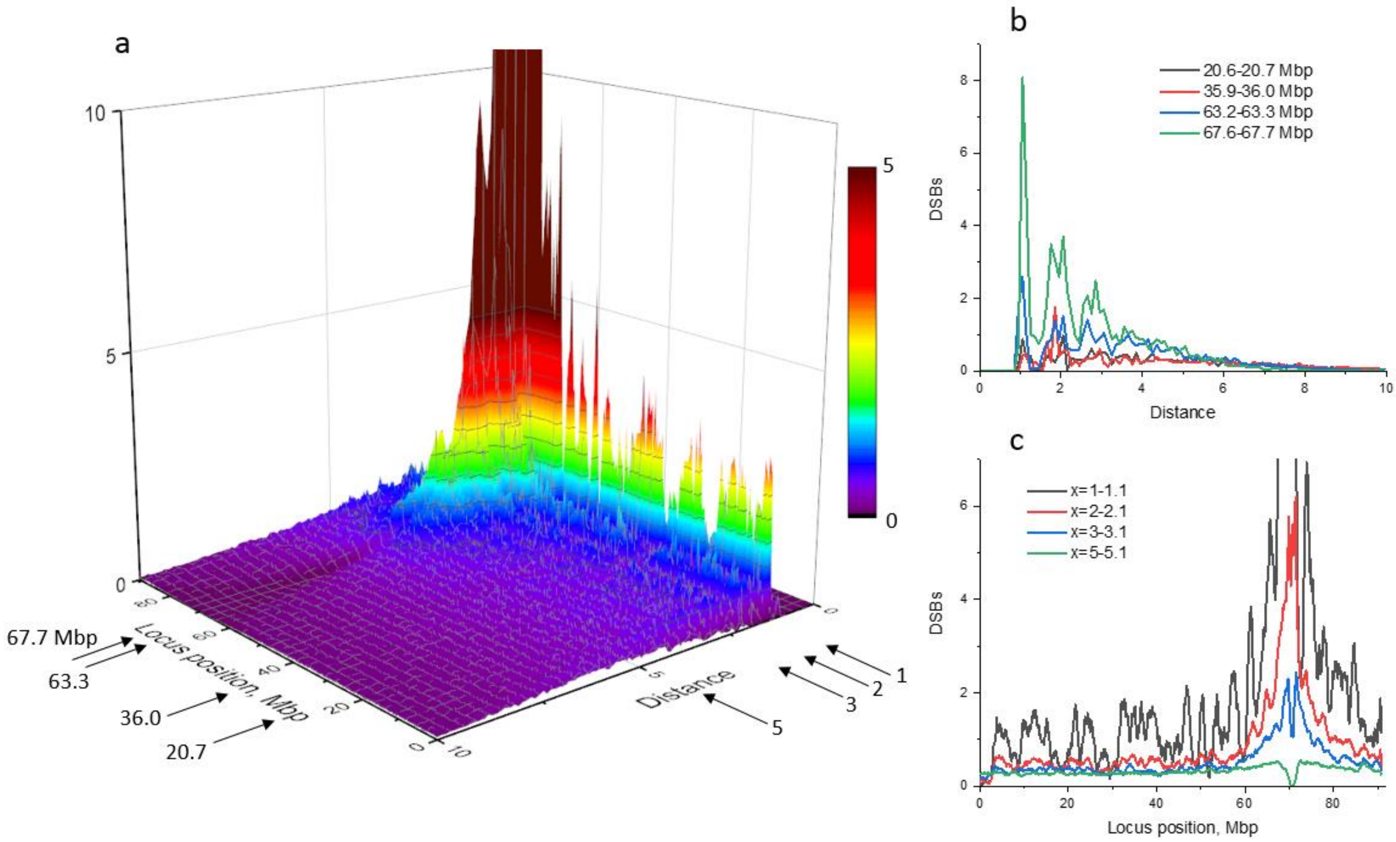


Figure S6. Complex interdependence between spatial and genomic separations for DSB distribution in chromosome 18. a: function $\varphi_{D,4C}(s,x)$ for D=5 Gy, heterogeneous globule, same as Fig.7 c. Arrows on the left indicate fixed positions of loci for $t_{4C}(x)$ calculations. Arrows on the right indicate fixed distances for $f_{4C}(s)$ calculations. b: spatial distributions $t_{4C}(x)$ for fixed locus positions. c: distributions $f_{4C}(s)$ for fixed distances.

### Supplementary Tables

Table S1. Characteristics of structures used

| | **Heterogeneous globule** | **Dense homogeneous globule** | **Loose homogeneous globule** | **Coil** |
|---|---|---|---|---|
| Mean gyration radius, µm | 1.11 | 0.99 | 1.06 | 8.40 |
| Contacts (s>1) | 2935 | 2730 | 1782 | 188.6 |

Table S2. Adjustable parameters of the model

| **Parameter meaning** | **Designation** | **Value** |
|---|---|---|
| Excluded volume potential coefficient | $U_{0ev}$ | 1kT |
| Initial attracting potential coefficient | $U_0$ | 1.2kT |
| Contact distance (between element centers) | $R_{cont}$ | 1.2 |
| Probability of incomplete exchange formation | $P_{inc}$ | 0 |
| Maximal distance of interaction | $R_{int}$ | 2 |
